# Capturing temporal heterogeneity of communities: a temporal β-diversity based on Hill numbers and time series analysis

**DOI:** 10.1101/2023.09.25.559413

**Authors:** Daniel J. Sánchez-Ochoa, Edgar J. González, María del Coro Arizmendi, Patricia Koleff, Raúl Martell-Dubois, Jorge A. Meave, Hibraim A. Pérez-Mendoza

## Abstract

Beta-diversity is a term used to refer to the heterogeneity in the composition of species through space or time. Despite a consensus on the advantages of measuring β-diversity using data on species abundances through Hill numbers, we still lack a measure of temporal β-diversity based on this framework. In this paper, we present the mathematical basis for a temporal β-diversity measure, based on both signal processing and Hill numbers theory through the partition of temporal ƴ-diversity. The proposed measure was tested in four hypothetical simulated communities with species varying in temporal concurrence and abundance and two empirical data sets. The values of each simulation reflected community heterogeneity and changes in abundance over time. In terms of ƴ-diversity, *q*-values are closely related to total richness (S) and show a negative exponential pattern when they increase. For α-diversity, *q*-value profiles were more variable than ƴ-diversity, and different decaying patterns in α-diversity can be observed among simulations. Temporal β-diversity shows different patterns, which are principally related to the rate of change between ƴ- and α-diversity. Our framework provides a direct and objective approach for comparing the heterogeneity of temporal community patterns; this measure can be interpreted as the effective number of completely different unique communities over the sampling period indicating either a larger variety of community structures or higher species heterogeneity through time. This method can be applied to any ecological community that has been monitored over time.

## Introduction

We live in a biodiverse world were species changes along space and time (1,2). The different forms of biodiversity have been studied using different mathematical, statistical and information system approaches (3,4). The principal goal of all measures is to characterize variation in biodiversity across different spatial and temporal scales (5–7). The regional component of biodiversity (γ diversity) contains the inter-site differences (β diversity) between local richness of species (α diversity). Thus, the study of γ, α and β diversity provides values that can predict changes on diversity (8–13). γ, α and β diversities are thus a group of key concepts for understanding the changes in species over landscape, and their applications are wide and include species dispersion, hotspot regionalization, reserve design, and clarification of the complementarity of the composition of species; it is thus one of the most important concepts of ecological theory (2,3,13–19).

The concepts of γ, α and β diversity revolves around the understanding of biodiversity patterns. In general, γ and α diversity are different mathematically and similar conceptually but not in scale, being α diversity focused at local and γ at regional. Otherwise, the concept of β-diversity revolves around the heterogeneity of species. The concept of γ diversity is simply the total number of species on a region. α diversity measures attach various definitions influenced by different assumptions of the species diversity, in general there are 2 main analytical groups, the ones related with absence presence data and the based abundance data. Contrary, β diversity measures use different mathematical contexts, concepts and frameworks. Nevertheless, there are two main analytical approaches for measuring β-diversity. The first one is the decomposition approach, in which β-diversity is calculated based on the segregation of ƴ-diversity and α-diversity components; the second one includes differentiation or variance measures derived from the total similarity of a pairwise community abundance matrix (20). Both approaches allow different hypotheses related to the processes driving species distribution and diversity patterns to be tested (3,13,20–22).

Differentiation measures are based on similarity-dissimilarity analysis, and the central idea is to assess changes in community structure from a sampling unit to another across different spatial, environmental, or temporal scales (23–25). Therefore, the differentiation approach can be used to address questions related to gains or losses of species among spatial or temporal scales and changes in their proportion among sites (23,26). In general, these last approaches involve community turnover and nestedness models, most of which are based on pairwise differences in space, time, or the environment (13).

The decomposition β-diversity approach is based on the overall component of diversity (ƴ- diversity) and its relationship to within-community local diversities (α-diversity). Under this approach, raw data are not directly used because the decomposition approach depends on the calculation of α-diversity, except when α = S. In general, this approach captures community variation for an entire set of sampling units that do not require exact species identities because it is based on the calculation of α-diversity (26,27). Consequently, this approach can be used to address questions related to the heterogeneity and the number of unique species in the communities in a landscape, or to the proportion of species that are not shared among all sampling units. Several multiplicative and additive algorithms can be used under this approach, but all of them are based on the relationship between α- and ƴ-diversities (27–29). Although there is much discussion about what measure is optimal for a given objective or purpose, it has been suggested that both approaches could provide complementary insights that contribute to the advancement of community theory (26,30). However, the temporal axis of β-diversity has been less explored than the spatial axis, and even some authors have pointed out the lack of reliable temporal frameworks and measures (3,31).

The approaches and frameworks for measuring temporal γ, and α diversity are related with the common spatial measures as Simpson, Shannon and Jaccard index (Shannon, 1948; Simpson, 1949; Jost, 2006b; Chung, Miasojedow, Startek, & Gambin, 2019). There is a unique temporal α diversity approach based on Hill numbers theory (Chao et al., 2021). This measure is the most robust and counts for the effective number of equally abundant species, and can also be applied for species traits (Chao et al., 2021). However, β-diversity measures are diverse and framework development is still lacking (31). Engen proposed to calculate bivariate correlations among community assemblages (24). Later, Baselga proposed the β_TUR_ and β_NES_ measures. These indexes are the most used temporal β diversity measures because they permit the use of presence/absence data, information that can be easily obtained with low sampling effort (13).

Legendre used Morańs Eigenvector Map analysis (MEMs) to calculate the positive and negative correlations among time points in community samples (32). Finally, the temporal β-diversity index was proposed to measure changes in species composition between adjacent time points (33). In general, all the proposed measures are similar to spatial diversity measures. These measures of temporal α and β-diversity are accessible, but the outputs of these measures are not completely comparable between studies, with exception with Chao et al (2021). The temporal β-diversity measures are more related to correlation measures, and the species composition among time points has been calculated using various approaches. Although all measures are quite robust to different traits and approaches, they are not immune to many of the problems associated with spatial β diversity measures. The main shortcomings associated with these measures include their tendency to generate misunderstandings of temporal traits and overlook data independence assumptions. Generally, most temporal biodiversity measurements are based on a sequence of data points indexed along a temporal axis. The data points are literally consecutive measurements of biodiversity of the same area over a time interval, which permits changes to be tracked over time. The assembly of biodiversity observations obtained by repeated measurements over time has certain peculiarities (34–36). Time series are normally characterized with information gaps created by sampling effort, which usually results in discrete variables. The amount of missing information will be determined by the time intervals between samples (34,37). Ecological data can present both stationary and non-stationary dynamics over time (XXX). Stationarity in a temporal context refers to the consistency in the mean and variance throughout time, thus a standardization of data should be performed. Any approach for estimating diversity from a temporal perspective has been developed without consideration of the aforementioned assumptions (33,38). Temporal diversity analysis has been conducted based on time-correlation analysis and time-to-time relatedness analysis of communities; consequently, comparisons among current studies of temporal diversity are not possible (24), except for α diversity (39). Thus, a robust temporal β-diversity analysis should be consider the above-mentioned time series assumptions relating to the gaps in the database (34,40,41). The major problem with this assumption is that ecological processes are considered to occur discretely, which is not true, moreover, species into a community present variation in their abundance changes. This assumption are effectively accounted through wavelet transform analysis, which includes the frequency estimators and provide a reliable approach for predicting gaps in the data to model a continuous variable (42). Finally, the standardization of the abundance time series data should be considered to account for the diversity in the temporal activity patterns of species over time (31). In light of these considerations, Hill numbers diversity theory integrated with time series analysis might provide a robust approach for overcoming the lack of comparability in current temporal measures. Hill numbers are a group of diversity metrics that have various advantages in the interpretation of diversity values principally because their units are consistent, which makes them duplicable and comparable among studies.

Hill numbers are a group of parametric measures used to measure diversity based on the modification of the *q* parameter (20,27), which determines the sensitivity of the measure to the relative abundance of species and can take any value ≥ 0 (27). Hill numbers denote the effective number of species in the community and have been shown to be reliable for characterizing several community traits (e.g., taxonomic, functional and phylogenetic diversity; Liebhold, Koenig, & Bjørnstad, 2004). The advantage of the Hill numbers approach is that results derived from it usage are more robust, comparable, and interpretable measures compared to other diversity index approaches because Hill numbers are direct diversity values (15,23,26,44).

A measure of temporal diversity based on Hill numbers and time series analysis that captures the heterogeneity of diversity through time could provide a more promising strategy for assessing this important biodiversity component, similar to the way in which spatial diversity measures have become comparable among studies (8,27). We expect that these measures will enhance estimates of temporal γ and α diversity, and have a new temporal β that allows interpretable comparisons among studies, considering the ecological processes as a continuous variable. In addition, the measures could be used to characterize temporal diversity patterns and thus provide insights into the temporal dynamics of communities; information on temporal compositional shifts can shed on light on the status of communities and the effects of environmental variables on temporal species heterogeneity (29,31,33).

## Materials and Methods

### Temporal patterns as a continuous variable: temporal diversity data preparation

We developed our temporal α, γ and β-diversity measures based on the wavelet transform analysis (45–47) and the Hill numbers diversity approach (26,27,48). Our proposed temporal diversity measures were developed specifically for the decomposition approach. By considering the fully stationary behavior of abundance species data and the gaps associated with variation in sampling effort, wavelet time series analysis was used to standardize and fill gaps in abundance information through an estimated continual abundance curve, and this permits the distance-based Hill numbers diversity measure to be used (26), with the area under the curve as the basis of the pairwise distance measure. Wavelet transform analyze the frequency spectrum of a discrete time series data, fitting a modeled smoothed curve. The analysis compares (through convolutions) the raw data with a scaling function which can be shrunk, stretched and shifted in time. Finally, a matrix is constructed with all the fitting values in which the sum of the columns results on the new modeled curve (Figure 1).

**Figure 1.**
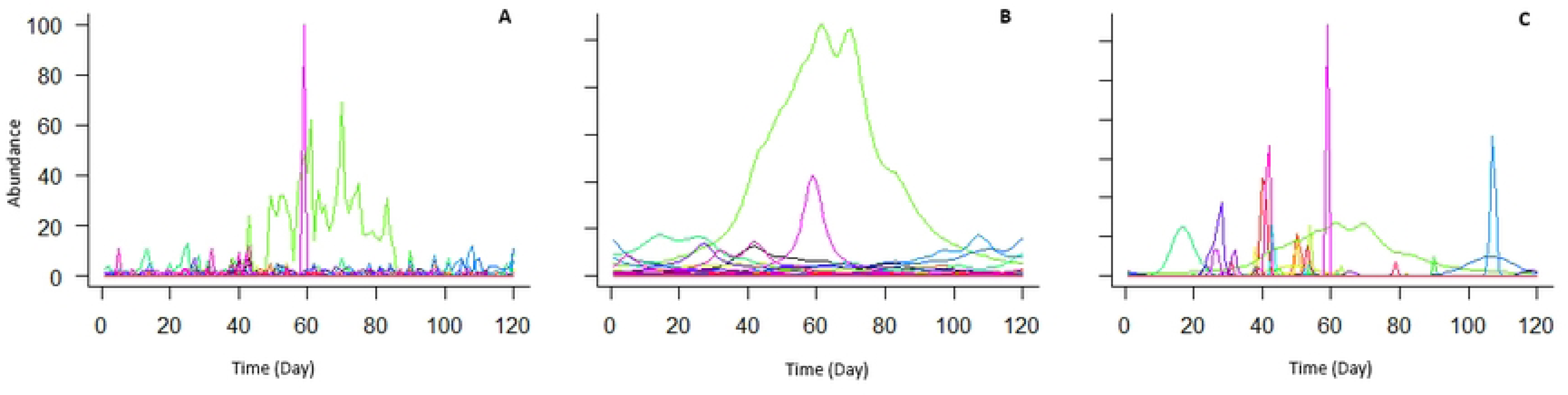
Comparison of same data set of abundance patterns of a Malagasy amphibian community. (A) representation of the raw data, (B) wavelet transformed data with τ = 2, and (C) wavelet transformed data with adjusted τ values. Each color represents a species (S = 40; N = 120 days).

### Validity of the stationarity of the time series of species abundance

The verification of stationary is pivotal for the standardization of the species abundance time series data. Generally, ecological data exhibit variability in their stationarity characteristics, often resulting in a limited degree of comparability when working with time series. For instance, we performed a stationarity test analysis to know the nature of species abundance curves data. We conducted an analysis to determine whether the temporal data need to be standardized through time series analysis.

### τ – the rate of change of species abundances over continuous time

Biological processes occur gradually and continuously, however in most cases they are discreetly recorded due to sampling. Nevertheless, there are differences in rates of change in the occurrence and abundance changes of species. For example, groups of species that have a high response to environmental changes often have abrupt changes in their abundance (such as amphibians, reptiles and insects). On the contrary, there are organisms in which changes in abundance occur in a more attenuated pattern (such as plant communities). Being more specific, within these generalities, it is true that each species has its particularities of the temporal variation, thus, τ is a variable that considers this biological aspect (Figure 1). Wavelet analysis typically assigns a value of 2 to the rates of change in modeled phenomena through time. Given that rates of change in the abundance of species through time are not equivalent due to variation in the traits of species, unique values of this parameter should be assigned to each species to account for this variation.

Wavelet analysis allows interpolation of a continuously modelled abundance curve; however, unlike the common use of wavelet analysis, this analysis for our purpose requires the specification of the attenuation threshold (τ), a parameter that controls for the rate of continuous change in the occurrence of species in the community. In other words, τ determines the slope (i.e., the speed) at which species abundance or species intensity changes through time (41,49). For example, there are communities where some species display faster rates of appearance with respect to others (50), i.e., depending on its intrinsic natural history traits, each species can potentially exhibit their own rate of change in abundance through time (51). Thus, a fixed parameter value (usually 2 in most analyses) is not an appropriate assumption, and a more objective value is needed for each species (47,52,53). In this way, a correct fixed τ must consider the resolution (sampling time interval), the total sampling effort (T), and the steepness of the abundance changes in the raw data. In this regard, the rate of change in the abundance of species over time and the accuracy of the data sampled are important. If time series data have a high resolution, τ will be weighted highly according to the rate of change in the abundance of each species; otherwise, when time series data have low resolution, τ-values will highly weight the new abundance curve calculated with wavelet transform analysis. Here, we propose, for a given *i* species, a value of

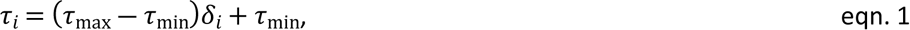

Where

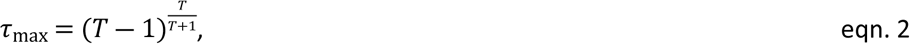

τ_max_ corresponds to a constant of the maximum value that τ_*i*_ can take in the analyzed community, τ_min_ = 2, τ_min_corresponds to the constant minimum threshold that τ_*i*_ can take. The equation

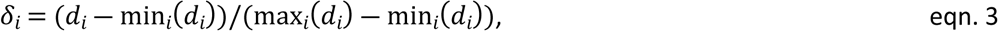

represents the rescaling factor between 0 -1 of *d*_*i*_, or the derivative that biologically refers to the rate of change where

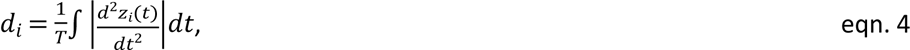

*d*_*i*_corresponds to the mean concavity of absolute values of the species abundance; in other words, *d*_*i*_ is the average steepness of the changes in species abundances between consecutive samples in time. Finally, *z*_*i*_(*t*) represents the raw abundance of the *i*th-species for each time, whereas *T* represents the total number of sampling times.

Each τ_*i*_ value integrates the rate of abundance change of each species based on the observations and the maximum number of sampling points of the study. The scaling function used for the wavelet analysis was the Morlet scaling function, which is optimal for data with unknown frequencies and scales, and data that cannot be directly interpreted (41).

### Temporal *β* diversity: effective number of distinct communities over time

For our temporal β-diversity measure, we replaced the discrete abundance vectors by abundance or intensity curves derived from the wavelet time series analysis. We modified the equations from Chao & Chiu (2014) (eqn. 5), so that the relative abundance values (*z_i_*) represent the abundance curves of the *i*-th species. To estimate temporal β-diversity through the multiplicative component (ƴ/α), we first need to calculate temporal ƴ-diversity and temporal α-diversity. To calculate ƴ- diversity of order *q* (*^q^DTγ*), we used the relative abundance of species in the community (*z*_*i*+_*/z*_++_; *i* = 1, 2, …, *S*), where *z*_*i*+_ = ∫_*T*_ *Z*_*i*_(*t*)*dt* is the total area of abundance of *i-th* species, and *z*_++_ = 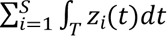 is the total sum of the area abundance of all species. Consequently, ƴ-diversity of order *q* is defined as

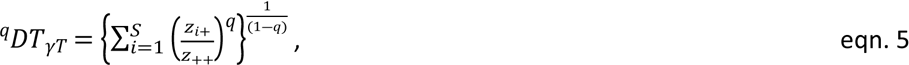

and when *q* = 1 as

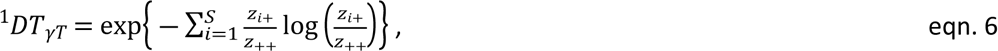

The temporal ƴ-diversity is interpreted as the “effective number of species in the entire community through time” or the species richness when q > 0. For temporal α-diversity we applied the same set of measures and definitions proposed by Chao & Chiu (2014) but on a temporal scale. In this sense, temporal α-diversity represents “the effective number of species per time unit” or the “mean effective number of species per time unit”, and defined by:

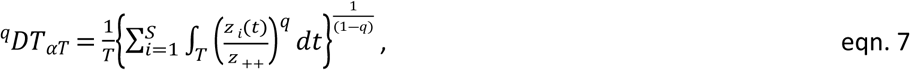

and when *q* = 1, as:

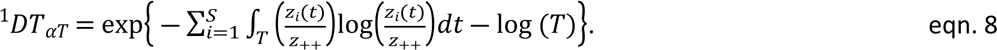

Finally, the multiplicative temporal β-diversity can be calculated as:

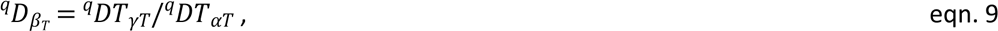

This value can be interpreted as “effective number of completely different unique communities over the sampling period”. The contribution of species heterogeneity among communities is based on changes in the rate of whole community richness and the mean community changes at each sampling point. Temporal α- and ƴ-diversities always range from > 0 to *S* and decrease as *q* increases; temporal β-diversity ranges from 1 (when *q* = 0) to infinite.

To illustrate the use and utility of this measure of temporal β-diversity, we performed simulations varying the abundance and heterogeneity of species richness. In addition, we performed two extra analyses based on field data of an amphibian community from Madagascar (*S* = 40; time period = 360 days; frequency = daily; Heinermann et al., 2015) and a macro-benthic community from Chesapeake Bay (*S* = 66; *time period* = 24 years, frequency = yearly; Chesapeake Bay Foundation, 2020).

We used R Studio with the DescTools (Asem, 2020) and wavScalogram (Benítez, Bolós & Ramírez, 2010) packages. The script for the temporal β-diversity calculation can be found in Appendix 1.

## Results

Patterns of temporal α- and ƴ-diversity were similar to those suggested by other diversity measures based on Hill numbers, which was consistent with expectation. The values of each simulation reflected community heterogeneity and changes in abundance over time. In terms of ƴ- diversity, *q*-values are closely related to total richness (S) and show a negative exponential pattern when they increase, except when species abundances are constant over time. For α-diversity, *q*- values profiles are more variable than for ƴ-diversity, and different decaying patterns in α-diversity can be observed among simulations (Figure 1). Whenever *q* increases, if there are differences in abundance, α-diversity values display a decaying pattern (Figures 1B, 1D and 1H), causing temporal β-diversity to show different patterns, as observed in the analyses based both on simulations and field data (Figures 1 and 2). Otherwise, the absence of a decreasing pattern in Figures 1 and 3 reflects the null variation in both species richness and abundances in the simulated data. The minimum values of β-diversity are always 1 and in general they increase as *q* increases, except in those cases where ƴ- and α- diversities do not change (i.e., Figure 1F and 1H). The mostvariable β-diversity pattern is shown in Figures 1B and 1D, where species are equally distributed over time, which corresponds to the most heterogeneous community in terms of species changes over time. For the Malagasy amphibian community, temporal β-diversity shows high values near *q* = 0 and then stabilizes as *q* increases; by contrast, β-diversity shows a nearly constant increasing pattern as *q* increases in the Chesapeake Bay macro-benthic community.

**Figure 2.**
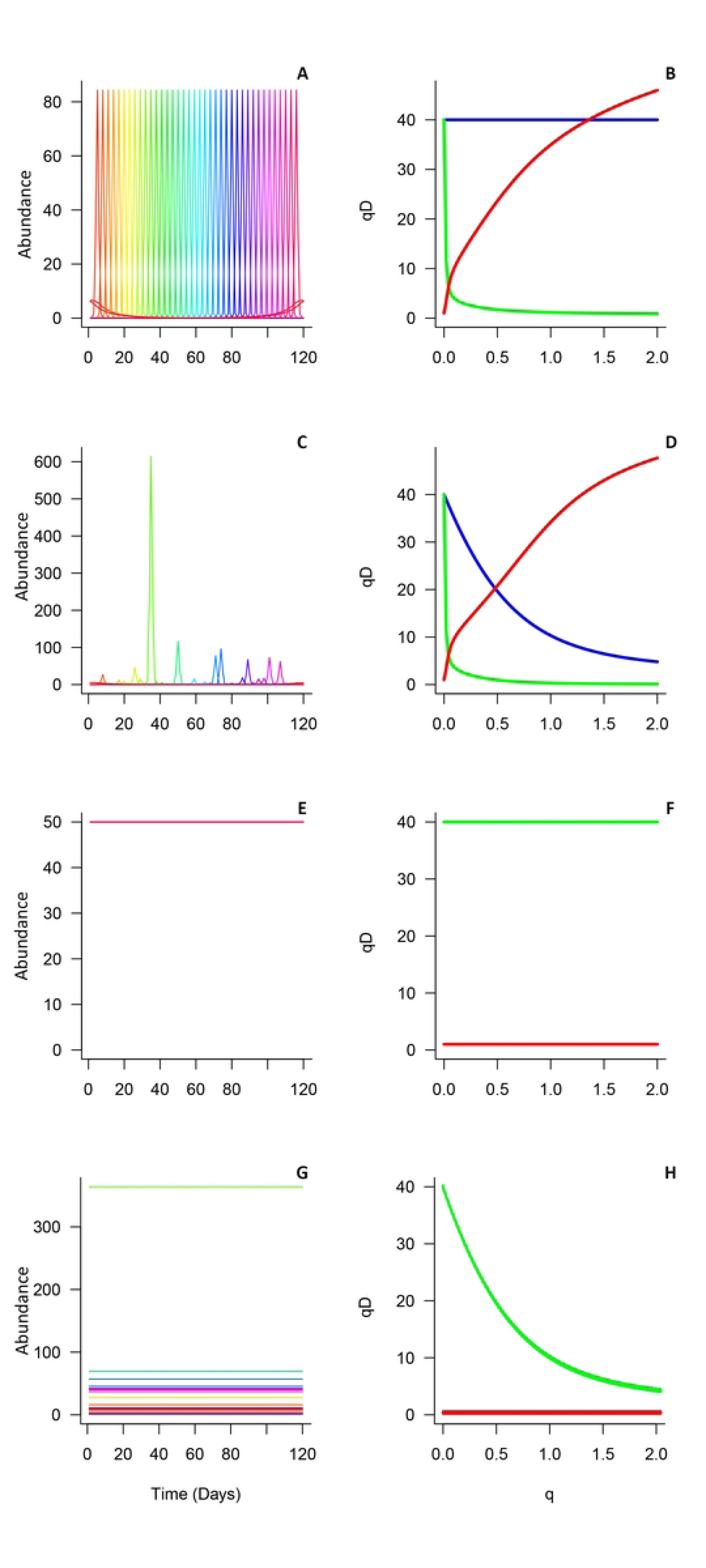
Graphs showing temporal ƴ-, α-, and β-diversity profiles as functions of q (0 ≤ *q* ≤ 2): (A) simulated data where species are equally distributed over time and abundance patterns are identical, representing a situation where each species can be found at any specific time. (B) Temporal ƴ- (blue line), α- (green line) and β-diversity (red line) profiles of panel A. (C) Simulated data where species are equally distributed over time and abundances are unequal. (D) Temporal ƴ- (blue line), α- (green line) and β-diversity (red line) profiles of panel C. (E) Simulated data where species are present all the time without abundance variation and equal abundances. (F) Temporal ƴ- (blue line), α- (green line) and β-diversity (red line) profiles of panel E. (G) Simulated data where species are present all the time without abundance variation and unequal abundances. (H) Temporal ƴ- (blue line), α- (green line) and β-diversity (red line) profiles of panel G.

**Figure 3.**
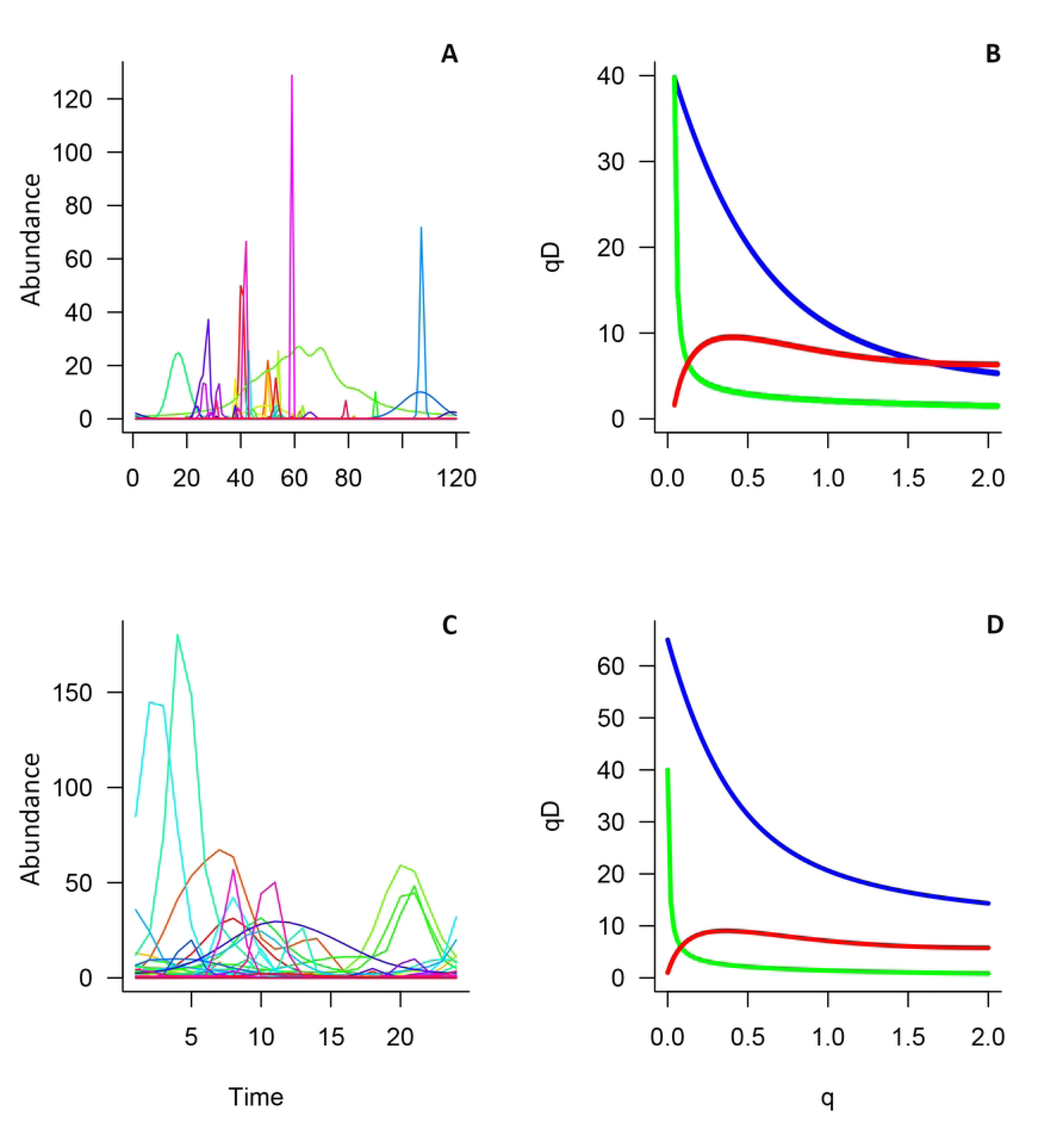
Graphs showing the temporal ƴ-, α- and β-diversity profiles as functions of q (0 ≤ *q* ≤ 2) for an amphibian community from Madagascar. (A) Wavelet transform distribution of species throughout time for the amphibian community from Madagascar (S = 40). (B) Temporal ƴ- (blue line), α- (green line) and β-diversity (red line) profiles of panel A. (C) Wavelet transform distribution of species throughout time for the macro benthos community from Chesapeake Bay (S = 66). (D) Temporal ƴ- (blue line), α- (green line) and β-diversity (red line) profiles of panel C.

## Discussion

Here, we propose a temporal γ, α and β-diversity measures based on time series analysis and Hill numbers diversity theory that provides an interpretable, efficient and comparable tool to characterize temporal community heterogeneity. The use of wavelet analysis for developing a temporal diversity framework enhanced the robustness of the results and circumvents sampling effort-related problems (42,47). To perform this analysis, we approximate the abundance time series data to simulate a continuous abundance curve as in other studies (37,49,52,54). One of the major challenges associated with temporal diversity analysis, especially β-diversity measures, is the failure of datasets to meet the assumptions of time series data because current measures do not consider sampling resolution patterns, time frequency characteristics (33,38,55), and stationarity of the variables; wavelet analysis can overcome this difficulty. Other studies have shown that time series analysis is an effective way to analyze data collected from ecological and forestry studies (52,56) for several purposes such as population dynamics, disease transmission, animal migration and phenology (45,49,52,57–60). Considering the great diversity in the activity patterns of species, we tested the stationarity in species abundance of our species to justify the use of wavelet analysis, given that wavelet analysis standardizes a continuously modeled curve, which permits comparisons of our beta-temporal diversity measure to be made. We suggest that the stationarity of species abundance should be tested as a best practice, as this provides key information for determining how the data could be optimally standardized. For our purpose, wavelet analysis provides an effective method for modelling a continuous abundance curve under the assumption that species abundance changes occur gradually rather than abruptly at different scales. Nevertheless, there are two assumptions to considerate in wavelet analysis transform related with sampling effort: Overall, (1) equidistant temporal sampling and (2) long time series data conduct to better modeled curves. It is possible to analyze data based on repeatedly sampling effort, depending on the question and life cycle of organisms, otherwise other time series analysis perspectives as Hilbert Huang approach should be better, but this perspective has not been used for ecological data. Likewise, is recommended that 25 is the minimum number of sampling temporal points necessary to wavelet analysis, nevertheless here we performed an example with 24 sampling points (Chesapeake microbenthic community) (37,61). There is no systematic assessment for that assumption and there is a need to test it with high resolution data, removing and reducing random sampling points.

Temporal changes in the abundance, conspicuousness or biological processes of species vary among all species in communities. Wavelet analysis through the *τ* parameter partially takes into account this variation. Mathematically, the *τ* parameter controls the slope of the modeled curve, but this parameter is not modified in conventional wavelet analysis (47,49,53). Biologically, the *τ* parameter reflects the rate of change in the occurrence and abundance of species; this prompted us to assign an objective *τ-*value for each species in the studied community. In this way, the *τ-* parameter directly relates to the rate of change in the abundance of each species and the total sampling points. In our study, *τ*-values near 2 imply that the analyzed species have abundance patterns similar to those modeled by wavelet analysis, and species with higher *τ*-values have simulated abundance curves that resemble the mode (statistics) of the raw data. Likewise, this idea suggests that the probability of detection or occurrence of species is a parameter that should be considered in all diversity studies, and the recorded frequencies of species are an artifact of specieś actual frequencies and their detection probabilities. The imperfect detection of species is important to consider to obtain high quality estimates of the size of ecological populations (62–64). This stems from the fact that the signals of species are underestimated when they are present but not detected. Various situations can lead to underestimates in the signals of species (37,49,65), and a correction in the detection in the abundance curves in our analysis could refine the overlapping area and τ calculation; nevertheless, imperfect detectability also varies with time.

For temporal α diversity, was possible to use the perspective of Chao 2016 (26), however it is not the same as that proposed in 2021 (39). Firstly, the already existing measure accounts for the effective number of equally abundant species and here, temporal α diversity accounts to the mean effective number of species seen per sampling unit. The differences between measures are mainly associated with the model construction, however, we considered the 2016 Chao’s perspective because it was a framework built to bridge the two β diversities approaches. On the other hand, this perspective indirectly considers the detection of organisms since it considers the average number of species seen per sample. Thus, this perspective is more sensitive to average changes in the number of species from sample to sample. Thus, this perspective is more sensitive to average changes in the number of species from sample temporal points. At the same time temporal α diversity values when q = 0 will be equal to those of temporal γ diversity when q = 0. This is mainly because the calculation of the continuous curve of abundance through wavelet assigns a value close to zero in the probability of encountering the species over time. In this sense, in this measure we are assuming that the species have imperfect detectability, and could be registered at any time, variating in four axes (species traits, spatial variation, temporal variation and sampling characteristics; (66). This has received a lot of attention at population and community level (50,64,67,68); however, some studies have pointed out the need to include imperfect detectability on biodiversity surveys (68,69). Future research on the standardization of values is needed to improve the comparability of results, without excluding low detectability or rare species. From our experience, link function, τ parameter, or even a cutoff value could be options where the diversity values could be better adjusted, however, with the change of these variables, diversity patterns remain unchanged.

The temporal β-diversity measure proposed here is a decomposition Hill numbers approach adopted from the scheme developed by Chao and Chiu (2014), which is based on the relationship between ƴ- and α-diversities in a community. Given the great variation in β-diversity decomposition methods, other decomposition approximations could also be tested using the same approach; our ƴ- and α-diversity calculations would be the same as the theoretical framework and can be easily linked to other perspectives (29,70). β-diversity can be analyzed through a distance-based approach; however, the Chao and Chiu framework was mainly constructed to establish a link between both β-diversity approaches (26) and a temporal β- distance-based approach to complement our measure, including time series transformation of the same data (15,71). Likewise, other temporal β diversity frameworks, such as those of Baselga and Legendre (13,33), have demonstrated the value of using multiple measures of β diversity, but the utility of using multiple measures ultimately depends on the hypothesis tested (9). However, the interpretation of these other measures requires caution because some refer to the concepts of turnover and others to variation (as our measure), but the interpretability is maintained under the Hill numbers framework. Otherwise, the most used measures of β diversity (Jaccard and Sorensen) (Jaccard, 1912; Sorensen, 1948; González, n.d.; Chao, Chazdon, & Shen, 2005; Baselga, 2012) and even α diversity (Shannon and Simpson) (74) do not have a unifying structure that facilitates interpretation and comparison (12,22).

Thus, the principal advantage of our framework is that the proposed temporal β-diversity measure provides a more direct and objective approach for comparing the heterogeneity of temporal community patterns. In this context, temporal ƴ-diversity is defined as the “effective number of species throughout the entire studied time period”, temporal α-diversity as the “mean effective number of species at each time”, and temporal β-diversity as the “effective number of completely different unique communities over the sampling period”. In general, ƴ- and α- *q* profiles are consistent across estimated spatial diversity patterns (44); however, *q*-profiles related to β- diversity do not show a consistent pattern. Specifically, temporal α-diversity only reflects an expected outcome rather than the reality indicated by the sampling measurements; thus, a completeness analysis could improve the robustness of the results for both temporal ƴ- and α- diversity as has been shown in other studies of diversity patterns (75–77). For temporal β- diversity, overestimations were observed in simulations, especially in cases where several species were equally distributed, as temporal β-diversity values are higher than *S* (the number of species) when *q* = 2. A high temporal β-diversity indicates a high number of unique communities throughout the sampling period and thus temporal heterogeneity in the activity of species within the community. Nevertheless, our measure is not suitable for indicating the moments where unique communities are occurring, but other measures can be used to provide this information, such as Legendre’s TBI (Temporal Beta Index) (33). Thus, we show here that the use of different frameworks provides complementary information and that the use of each measure is not mutually exclusive.

Despite differences in the taxonomic group, species richness, and temporal resolution among field studies, temporal β-diversity can be measured using these data. Although we expected asymptotic behaviors, we observed different temporal β-diversity patterns in the two data sets examined. In the Malagasy amphibian community, we observed that the temporal β-diversity *q*-profile shows high values when *q* is between 0 and 1, and the profile shows an asymptotic pattern. The rate of change in the α-diversity *q*-profile largely determines the heterogeneity of the community (temporal β-diversity) because α- and ƴ-diversity values are divergent. For example, few species of amphibians are commonly observed per sampling occasion or per unit of time; in other words, few species are recorded during each sampling event, and the species observed continuously vary as has been documented in other studies (78–80). This result has direct implications for our understanding of the heterogeneity of communities through time as well as for conservation and monitoring actions because some community traits through spatial and temporal scales exhibit divergent patterns (81,82). Thus, to understand the temporal relations as spatial perspectives of γ, α and β diversity is to analyze several data sets. It is true that the relation between spatial γ and α diversities largely determines spatial β diversity, something that we also suspect occurs in temporal diversity. From a temporal view, probably there is a linear correlation between temporal γ and α diversity, resulting in a constant and low temporal β diversity values. Thus if temporal α diversity (mean number of species per time) present low values per unit of time, temporal β diversity should be higher, resulting a temporarily diverse community (3,83). Communities that show high temporal heterogeneity in composition require conservation or monitoring plans in which sampling effort is high in frequency, so that the range of environmental variation that can occur at a site is sampled; however, the reason for the need for a high sampling frequency is not solely because of temporal variation in the composition of communities but also because of variation in the detectability of species as aforementioned (84). However, low temporal α-diversity values do not prove that some species do not occur in the studied unit; rather, it is likely that features of the environment affect their conspicuousness as several studies have shown (62,79,80) or even undergo short-distance migrations (85–87). All of these assumptions directly relate to other ecological processes such as interactions and phenological patterns because the occurrence of some species depends directly on the presence of other species (63); for example, the common interactions between flowering plants and pollinator life cycles (88,89), as well as interactions between predators prey (90–92) or parasites and host (93). In this way, patterns of temporal β diversity between different functional groups require comparison to determine whether the temporal β diversity of one group predicts the temporal β diversity of another group; our measure permits these comparisons to be made and other hypotheses to be tested. Finally, in the case of the Chesapeake Bay macro-benthic community, we observed that the *q*-profile of the temporal β- diversity increases without reaching an asymptote; thus, temporal β-diversity values are likely higher than the one presented (8.17) and a higher sampling resolution or a longer time window could alter these results; ultimately, it is likely that this community is more heterogeneous through time. This demonstrates the need for more studies that estimate temporal β diversity using different levels of sampling effort or conducting analyses at different time scales to understand the effect of scale on temporal β diversity patterns, as other studies have shown that scale affects diversity patterns in other ecological axes (94–96).

The implementation of our new temporal-diversity measure is needed to advance our understanding of community temporal species changes and its heterogeneity and how this could become a tool for the optimization of time and resources in management plans and community monitoring programs. Otherwise, understanding of temporal ecological patterns and their relationships with environmental cues could generate new questions related to temporal community changes and how communities are affected by this poorly explored axis. It would also be interesting to know whether temporal β-diversity responds similarly to spatial measures of temporal α- and γ-diversities. Finally, we emphasize that the temporal diversity measure proposed here is suitable for the analysis of any taxonomic level, community and temporal scale. Therefore, if we have a long time series data of species abundance, we are able to compare between years, seasons or any periodical perspective; always taking into account the wavelet limitations aforementioned or our own sampling resolution. Finally, this proposal looks for establish a baseline (principally for β diversity), for analyze temporal diversity.

## Conclusions

The analysis of temporal diversity is crucial for understanding the temporal distribution of species assemblages and the uniqueness and heterogeneity of species in communities. As the collection of long-term data increases, appropriate temporal analytical methods are needed to improve our understanding of temporal community patterns. Our temporal diversity framework produces intuitive, comparable and simple values for assessing species heterogeneity over time. Our measure has the same properties of other γ, α and β-diversity measures and can be applied to mid- and long-term community data sets available for any taxon even on disturbed ecosystems. Temporal α-, β- and ƴ-diversities have important implications for the temporal design of community monitoring, conservation and restoration programs.

## Acknowledgements

We are grateful to Miguel Vences for granting us access to the Madagascar amphibian community data. The first author acknowledges the Prosgrado en Ciencias Biológicas of Universidad Nacional Autónoma de México (UNAM) and the Mexican Science Council (Consejo Nacional de Ciencia y Tecnología, CONACYT) for a doctoral scholarship (2018- 000068-02NACF-21668). This paper is a partial requirement to obtain the Ph.D. degree.

## Author contributions

D.S-O., E.G., M.C.A, P.K., J.M and H.P. conceived the ideas and design of the study; D.S-O, E.G. and R.M-D. designed the methodological framework; D.S-O. obtained the data; D.S-O and E.G. analyzed the data; D.S-O, E.G. and H.P. wrote the manuscript. All authors contributed critically to the development of the ideas, approaches and draft, and gave final approval for publication.

## Data Availability

Malagasy amphibian community data available from the paper <u>10.1080/00222933.2015.1009513</u> (Heinermann et al. 2015) and Chesapeake Bay macro-benthic community data available from page https://www.chesapeakebay.net/what/downloads/baywide_benthic_database

